# Pre-departure psychological distress and associated factors among migrant workers of Nepal

**DOI:** 10.1101/2020.12.14.422661

**Authors:** Om Prakash Poudel, Bijay Thapa, Shital Bhandary

**Affiliations:** School of Public Health, Patan Academy of Health Sciences, Lalitpur Nepal

## Abstract

**Introduction:** Foreign employment is the most significant motivation for international migration in Nepal. However, migrant workers are vulnerable to many exploitations that lead to psychological distress during the pre-departure phase and at the destination. The study aimed to identify the prevalence and associated factors for psychological distress among migrant workers during the pre-departure phase.

**Methods:** This was a cross-sectional study based on the representative sample of 445 migrant workers. A 21-item Depression Anxiety Stress Scale (DASS-21) and Pre-Departure Risk Factors Perception Scale (PD-RFPS) at the workplace were self-administered to migrant workers selected randomly attending the pre-departure orientation program. Bivariate and multivariate logistic regression was performed to identify the associated factors.

**Results:** Prevalence of psychological distress (Depression, Anxiety and Stress) was identified as 20.9% and female (AOR=2.02, p-value=0.041) and perception of bad working conditions (AOR=2.44, p-value=0.046) were found significantly associated with pre-departure psychological distress.

**Conclusion:** Data suggests the presence of symptoms of psychological distress among migrant workers during the pre-departure phase and perception of risk factors at the workplace were found significantly associated with pre-departure psychological distress. Concern bodies should provide in-depth orientation on possible risk factors at the destination and coping skills for psychological distress during the pre-departure orientation program.

## Introduction

The number of international migration has continued to grow and estimated to around 272 million in 2019. Among these migrants, most of them were of the working age group (20-64 years of age) [1]. With the increase in the number of international migration, the global flow of remittance had been increased in the recent decades [2]. In Nepal, foreign employment is the most significant motivation for international migration. Although India is the popular destination for Nepalese migrant workers, it has been decreasing with the rise of labor migration in Gulf Cooperation Council (GCC) and Malaysia in the last decades. Limited opportunities, responsibilities towards family well-being, and attraction of Gulf countries are the major reasons behind the migration towards GCC countries [3]. In Nepal, more than half of the migrant workers acquired a permit to GCC countries and around thirty percent of permits were issued for Malaysia during past decades [4].

Migrant workers face different kinds of challenges in personnel, family, and social life. These workers go through different phases that include pre-departure, short-term and long-term transient, destination situation, and return to a place of origin. In each phase, potential health risk and protective factors exist that have a short-term or long-term impact on the well-being of migrant workers [5]. Most of the Nepalese migrants work in semi-skilled and low skilled jobs, which are often difficult, dangerous and degradable leading to poor mental health [6]. In Nepal, 607 deaths among male and 43 among female were reported due to suicide while working abroad, during the period of 2008/2009 to 2016/2017 [4].

Pre-departure and in-service at the destination are the two major stages where migrant workers are vulnerable to exploitation like manipulation of contract, hiring for non-existent jobs, poor working conditions, wages below standard, health and safety risk, and prolonged debt period that results in psychological distress [7]. Therefore, instead of focusing only on tertiary prevention, targeting the pre-departure determinant of the problem would be the most appropriate approach as primary prevention to improve the psychological well-being of the migrant workers. For that reason, the objective of this study was to identify the prevalence of psychological distress and its associated factors among migrant workers of Nepal departing to GCC countries.

## Methods

This was a cross-sectional study conducted among migrant workers attending a pre-departure orientation program. A total of 445 migrant workers were included from eight randomly selected orientation-training institutes out of 112 institutes inside Kathmandu valley from July to August 2019. Random allocation of the migrant worker in the study was ensured by selecting the week to visit institutes randomly. Thereafter, each randomly selected institutes were visited for one week and all the eligible migrant workers were approached in the study. Those migrant workers traveling to GCC countries were included in the selection criteria. Whereas, illiterate and non-Nepali speaking migrant workers were excluded. The questionnaire was self-administered to all the eligible consented migrant workers. Ethical approval was obtained from the Institutional Review Committee – Patan Academy of Health Sciences with reference PHP1906281263.

Socio-demographic and foreign employment related factors were taken as independent variables whereas psychological distress was taken as a dependent variable in the study. Sociodemographic variables included age, sex, religion, ethnicity, education, marital status, occupation, personnel behavior, and place of residence. Similarly, foreign employment related variables included pre-departure related factors and perception of migrant workers on risk factor at the workplace. Pre-departure related factors were frequency of travel, the reason for migration, loan, pre-arranged accommodation at the destination, pre-arranged employment at the destination, family network available at destination, the language proficiency of destination and discussion with family. Likewise, PD-RFPS [8] was used to measure the perception of risk factors in the workplace. Psychological distress was defined as combination of depression, anxiety and stress and it was measured using DASS-21 scale. DASS-21 is, a globally validated standard tool to measure depression, anxiety, stress and psychological distress. DASS-21 measures the severity of symptoms of depression, anxiety, and stress, containing seven questions for each subscale [9]. DASS-21 has already been validated in the Nepali language among Nepalese migrants residing in Hong Kong [10]. Besides, it has been validated among Nepali migrant workers during the pre-departure phase from the pilot study, which has been reported elsewhere [8].

Statistical analysis (Descriptive and analytical statistics) was performed using Stata 13 MP version software. The prevalence of psychological distress was calculated by adding the DASS total score, which indicates the overall index of negative effects [9]. The score was multiplied by the factor of two to make comparable with DASS-42. A score of more than 40 was included to calculate the prevalence of psychological distress indicating the symptoms of moderate or above in the DASS total score. Bivariate and multivariate logistic regression was performed to identify the associated factors for psychological distress. Variables found with the p-value of less than 0.25 in the bivariate logistic regression were included in the final multivariate logistic regression to see the independent effect of the variable on the symptoms of psychological distress.

## Results

Symptoms of psychological distress were experienced by 20.9% of migrant workers during the pre-departure phase. Symptoms of psychological distress were more prevalent among females compared to males. Nearly one-fourth of younger age groups less than 25 years were found with the symptoms of psychological distress, which was higher than the older age group migrant workers. Similarly, it was high among those who had smoked in the last month compared to those who had both smoked and consumed alcohol. Likewise, nearly one-third of unskilled manual workers were found with symptoms of psychological distress and it was less among the professional jobholders.

Psychological distress was high among those who were traveling for the second time compared to the first time travelers. Those who were traveling due to poverty, family pressure, and to pay back debt were found with a higher prevalence of symptoms. Similarly, it was high among those who had not discussed with their family about foreign employment, had a loan, had not prearranged for employment, had not pre-arranged accommodation, had no family member at the destination, and who did not understand the language of the destination country.

**Table 1:** Prevalence of psychological distress among migrant workers by independent variables (n=445)

| Variable | Number | Percentage (95% CI) | Variable | Number | Percentage (95% CI) |
| --- | --- | --- | --- | --- | --- |
| **Total** | 93 | 20.9 (17.1 – 24.7) | **Gender** |  |  |
| **Age Group** |  |  | Male | 71 | 19.6 (15.5 – 23.7) |
| Less than 25 | 45 | 24.6 (18.3 – 30.9) | Female | 22 | 27.2 (16.9 – 37.4) |
| 25-34 | 40 | 18.5 (13.2 – 23.8) | **Personnel Behavior** |  |  |
| 35-44 | 7 | 16.3 (5.4 – 27.2) | Alcohol only | 7 | 20.2 (7.7 – 32.6) |
| 45 or More | 1 | 23.1 (-15.0– 61.2) | Smoking only | 12 | 25 (12.7 – 37.2) |
| **Place of residence** |  |  | Both | 9 | 15.9 (6.3 – 25.4) |
| Urban | 46 | 18.4 (13.5 – 23.3) | None | 65 | 21.3 (16.6 – 25.9) |
| Rural | 47 | 23.9 (18 – 29.9) | **Frequency** |  |  |
| **Ethnicity** |  |  | First | 38 | 19 (13.4 – 24.5) |
| Brahman | 6 | 14.4 (3.3 – 25.5) | Twice | 42 | 24.9 (18.4 – 31.4) |
| Chettri | 24 | 23.4 (15.1 – 31.6) | More than two | 13 | 16.9 (8.6 – 25.2) |
| Newar | 4 | 16.3 (1.2 – 31.4) | **Reason for travel** |  |  |
| Janajati | 45 | 22.7 (16.8 – 28.6) | Family pressure | 3 | 30.7 (2.6 – 58.8) |
| Dalit | 10 | 20.5 (9.1 – 31.9) | Family conflict | 2 | 62.5 (2.4 – 122.5) |
| Muslim | 2 | 21.4 (-2.5 – 45.4) | lack of employment | 43 | 18.1 (13.2 – 23.1) |
| Madhesi | 2 | 8.6 (-3.9 – 21.3) | less paid at work | 11 | 13.4 (5.8 – 21) |
| **Religion** |  |  | Low agriculture production | 2 | 17.8 (-4.5 – 40.2) |
| Hindu | 72 | 22.2 (17.6 – 26.8) | To payback debt | 10 | 27.1 (12.2 – 42) |
| Buddhist | 15 | 17.4 (9.3 – 25.5) | Poverty/Landlessness | 19 | 33.1 (20.8 – 45.2) |
| Christian | 2 | 13.3 (-5.8 – 32.4) | Present political condition | 2 | 38.4 (-5.6 – 82.5) |
| Muslim | 3 | 22.2 (0.9 – 43.6) | Others | 1 | 15.3 (-17.3– 48.1) |
| Kirat | 1 | 14.2 (-9.7 – 38.2) | **Discussion with family** |  |  |
| **Marital Status** |  |  | Yes | 90 | 20.7 (16.9 – 24.5) |
| Unmarried | 36 | 21.2 (14.9 – 27.5) | No | 3 | 28.5 (-2.4 – 59.5) |
| Married | 56 | 20.7 (15.9 – 25.5) | **Take loan** |  |  |
| Widow/Widower | 1 | 40 (-48.1 – 128.1) | Yes | 74 | 22.8 (18.2 – 27.4) |
| **Education Level** |  |  | No | 19 | 15.7 (9.2 – 22.1) |
| Primary | 11 | 20.9 (9.2 – 32.5) | **Pre-arranged employment** |  |  |
| Secondary | 54 | 23.9 (18.3 – 29.4) | Yes | 76 | 20.4 (16.3 – 24.5) |
| Higher secondary | 22 | 16.7 (10.2 – 23.1) | No | 17 | 23.2 (13.4 – 32.9) |
| University | 6 | 17.2 (4.6 – 29.9) | **Pre-arranged accommodation** |  |  |
| **Employment** |  |  | Yes | 75 | 20.5 (16.3 – 24.7) |
| Un-employed | 38 | 22.6 (16.3 – 28.9) | No | 18 | 22.5 (13.4 – 31.7) |
| Agriculture | 26 | 26.6 (17.6 – 35.6) | **Family/Relative network** |  |  |
| Unskilled manual | 7 | 30.6 (10.6 – 50.6) | Yes | 63 | 19.4 (15.1 – 23.8) |
| Skilled manual | 13 | 15.1 (7.6 – 22.6) | No | 30 | 24.3 (16.8 – 31.7) |
| Business | 6 | 16.4 (4.3 – 28.5) | **Language** |  |  |
| Professional/Job | 3 | 8.6 (-1.5 – 18.9) | Yes | 36 | 18.4 (12.9 – 23.9) |
|  |  |  | No | 57 | 22.8 (17.6 – 28.0) |

CI = Confidence Interval

In bivariate logistic regression, none of the socio-demographic and foreign employment related variables were found associated with psychological distress except for the perception of a risk factor at the workplace. Perception of bad working condition (OR=4.28), not following contract properly (OR=3.21), poor safety measures at work (OR=3.15), not getting salary as contracted (OR=2.42), poor living place (OR=2.11), not getting leave (OR=1.99), food problem (OR=1.83) and not getting rest at work (OR=1.81) were significantly associated with psychological distress. Final multivariate analysis showed psychological distress significantly associated with the female (AOR=2.01) and perception of a bad working condition at work (AOR=2.44).

**Table 2:** Bivariate and multivariate logistic regression of psychological distress

| Variable | OR | 95% CI | p-value | AOR | 95% CI | p-value |
| --- | --- | --- | --- | --- | --- | --- |
| **Gender** |  |  |  |  |  |  |
| Male | Reference |  |  | Reference |  |  |
| Female | 1.52 | 0.86 – 2.71 | 0.145 | 2.02 | 1.03 – 3.94 | 0.041 |
| **Employment** |  |  |  |  |  |  |
| Unemployed | 3.07 | 0.82 – 11.43 | 0.094 | 3.50 | 0.75 – 16.37 | 0.112 |
| Agriculture | 3.81 | 0.99 – 14.62 | 0.051 | 3.65 | 0.74 – 17.97 | 0.112 |
| Unskilled manual | 4.63 | 0.97 – 22.08 | 0.054 | 2.77 | 0.44 – 17.27 | 0.275 |
| Skilled manual | 1.88 | 0.46 – 7.55 | 0.373 | 1.77 | 0.35 – 8.79 | 0.485 |
| Business | 2.07 | 0.44 – 9.57 | 0.352 | 2.04 | 0.35 – 11.99 | 0.431 |
| Professional/Job | Reference |  |  | Reference |  |  |
| Place of residence |  |  |  |  |  |  |
| Urban | Reference |  |  | Reference |  |  |
| Rural | 1.39 | 0.88 – 2.20 | 0.155 | 1.31 | 0.77 – 2.20 | 0.312 |
| Frequency of travel |  |  |  |  |  |  |
| First | 1.15 | 0.58 – 2.28 | 0.688 | 0.87 | 0.38 – 1.95 | 0.734 |
| Twice | 1.62 | 0.82 – 3.20 | 0.159 | 1.44 | 0.67 – 3.08 | 0.347 |
| More than two | Reference |  |  | Reference |  |  |
| Reason for Travel |  |  |  |  |  |  |
| Family pressure | 2.44 | 0.18 – 32.86 | 0.500 | 1.20 | 0.07 – 20.59 | 0.899 |
| Family conflict | 9.16 | 0.39 – 211.1 | 0.166 | 7.48 | 0.25 – 224.67 | 0.247 |
| Lack of employment | 1.22 | 0.12 – 12.18 | 0.863 | 0.61 | 0.05 – 7.17 | 0.698 |
| Less paid at work | 0.85 | 0.08 – 9.06 | 0.894 | 0.59 | 0.05 – 7.46 | 0.683 |
| Low agriculture production | 1.19 | 0.08 – 17.83 | 0.897 | 0.55 | 0.03 – 10.23 | 0.688 |
| To payback debt | 2.05 | 0.18 – 22.41 | 0.556 | 0.80 | 0.06 – 10.51 | 0.866 |
| Poverty | 2.71 | 0.26 – 28.19 | 0.402 | 1.45 | 0.12 – 17.44 | 0.771 |
| Present political condition | 3.43 | 0.20 – 58.33 | 0.393 | 4.45 | 0.21 – 93.36 | 0.336 |
| Others | Reference |  |  | Reference |  |  |
| Loan |  |  |  |  |  |  |
| Yes | 1.59 | 0.91 – 2.76 | 0.098 | 1.22 | 0.65 – 2.27 | 0.530 |
| No | Reference |  |  | Reference |  |  |
| Perception of risk factors at workplace |  |  |  |  |  |  |
| Bad working condition | 4.28 | 2.16 – 8.44 | 0.000 | 2.44 | 1.02 – 5.87 | 0.046 |
| Poor safety measure | 3.15 | 1.71 – 5.83 | 0.000 | 1.50 | 0.67 – 3.39 | 0.327 |
| Work other than told | 1.47 | 0.88 – 2.46 | 0.141 | 0.86 | 0.44 – 1.67 | 0.649 |
| Work long hour | 1.41 | 0.89 – 2.24 | 0.145 | 1.02 | 0.57 -1.83 | 0.942 |
| Festival problem | 1.38 | 0.87 – 2.20 | 0.164 | 1.09 | 0.59 – 1.99 | 0.780 |
| Food problem | 1.83 | 1.15 – 2.91 | 0.010 | 1.78 | 0.96 – 3.32 | 0.069 |
| Unfavorable weather | 1.36 | 0.85 – 2.18 | 0.194 | 1.22 | 0.69 – 2.14 | 0.484 |
| Did not follow contract | 3.21 | 1.79 – 5.78 | 0.000 | 2.03 | 0.92 – 4.36 | 0.080 |
| Cannot change work | 0.66 | 0.41 – 1.05 | 0.080 | 0.59 | 0.34 – 1.02 | 0.061 |
| Did not get salary as contracted | 2.42 | 1.39 – 4.19 | 0.002 | 1.63 | 0.75 – 3.56 | 0.221 |
| Did not get leave | 1.99 | 1.04 – 3.78 | 0.036 | 1.06 | 0.46 – 2.47 | 0.888 |
| did not get paid leave | 1.44 | 0.91 – 2.28 | 0.119 | 1.15 | 0.67 – 1.98 | 0.606 |
| Get unnecessary problem | 1.34 | 0.85 – 2.13 | 0.209 | 0.81 | 0.45 – 1.46 | 0.490 |
| Did not get rest | 1.81 | 1.12 – 2.94 | 0.016 | 1.35 | 0.74 – 2.46 | 0.331 |
| Poor living place | 2.11 | 1.17 – 3.82 | 0.013 | 0.89 | 0.41 – 1.96 | 0.783 |
| Cannot send money on time | 1.70 | 0.93 – 3.12 | 0.085 | 0.69 | 0.29 – 1.62 | 0.396 |

OR = Odds Ratio; AOR = Adjusted Odds Ratio; CI = Confidence Interval

## Discussion

The objective of this study was to identify the prevalence of psychological distress and its associated factors among migrant workers of Nepal departing to GCC countries. Although the psychological distress among Nepali migrant workers was studied at the destination and after return, there is a dearth of evidence regarding the mental health of migrant workers during the pre-departure phase. Psychological distress had been found significant among migrant workers during the pre-departure phase from this study. Besides, the perception of different risk factors at the workplace were found significantly associated with pre-departure psychological distress.

This study showed symptoms of psychological distress (DASS total score > 40) among 20.9% of migrant workers during the pre-departure phase. The prevalence of psychological distress in this study was found higher than the prevalence reported in recent studies conducted among returnee Nepalese migrant workers from India using the 12-item General Health Questionnaire (GHQ-12) tool to measure psychological distress [11,12]. A recent study conducted by the International Organization for Migration in Nepal had found psychological distress among 13.4% of returnee migrant workers from India [12]. Similarly, psychological distress was also reported in high number among Sri-Lankan aspiring migrant workers, which reported as 44.2% from GHQ-12 [13]. The difference in the stage of migration and tool to measure prevalence might be the reason for the higher prevalence of psychological distress.

Present findings are found similar to the reported mental health problem among migrant workers in different phases of migration in recent national and international studies [6,14]. Poor mental health was reported by 23% of returnee migrant workers in Nepal [6]. Likewise, a 22% of nondomestic migrant workers in Singapore were found at higher risk of psychological distress [14]. Psychological distress reported in this study was also found similar with the depression and anxiety among migrant workers reported in the systematic analysis of the studies conducted globally, which was 20% and 21% respectively [15].

From the multivariate analysis, females and those who had a perception of the bad working condition were found twice at risk of getting symptoms of psychological distress compared to males and those who had a perception of a good working condition at destination. Perception and post-migration expectations were associated with the occurrence of stress during the premigration phase [16]. A global review of literature on factor associated with common mental disorder among migrant population also reported that women and migrants who had bad working condition had higher mental health problems [17]. Similar to the present findings, Nepalese returnee migrants who had perceived risk and bad working condition at destination had also reported two times more mental health problems [6]. Other perceptions regarding working and living environment at destination were found associated with pre-departure psychological distress in bivariate analysis. These risk factors were also identified significantly associated with different health and mental health related problems among migrant workers in different studies [17–20]. Although Nepalese migrant workers are familiar with the risk associated with employment at Gulf countries, limited opportunities and responsibilities of a family at the place of origin are the reason behind simply accommodating these risks [3].

The limitation of the study was the exclusion of illiterate, non-Nepali speaking, and those who were traveling to the countries other than the GCC countries, which lacks the representation of all the migrant workers of Nepal.

## Conclusion

Findings from the study indicate that migrant workers face significant challenges for poor mental health in all phases of migration including the pre-departure. Different perceptions of work and living environment at destination were found significantly associated with pre-departure psychological distress indicating the need for in-depth orientation before departure to the destination. Since females were found more at risk of getting symptoms of psychological distress, further quantitative, as well as a qualitative study, are required to understand the causes of psychological distress during the pre-departure phase. As well, migrant workers could be provided with the coping skills for psychological distress during their pre-departure orientation program.

## Acknowledgments

We appreciate the support and feedback provided by all the faculty of the School of Public Health, Patan Academy of Health Sciences. We would like to thank all the participants who took part in the study.

## References

1. IOM. World Migration Report 2020. 2020. Available from: https://www.iom.int

2. IOM. World Migration Report 2018. 2018. Available from: https://www.iom.int/wmr/world-migration-report-2018

3. Binayak Malla MSR. Understanding Nepalese Labor Migration to Gulf Countries. J Poverty. 2017;21(5):411–33. doi: https://doi.org/10.1080/10875549.2016.1217578

4. Government of Nepal, Ministry of Labour, Employment and Social Security. Labour Migration for Employment: A Status Report for Nepal:2015/2016-2016/2017. Gov Nepal. 2018. Available from: https://asiafoundation.org/wp-content/uploads/2018/05/Nepal-Labor-Migration-status-report-2015-16-to-2016-17.pdf

5. Abubakar I, Aldridge RW, Devakumar D, Orcutt M, Burns R, Barreto ML, et al. The UCL-Lancet Commission on Migration and Health: the health of a world on the move. Lancet. 2018;392. doi: 10.1016/S0140-6736(18)32114-7

6. Adhikary P, Sheppard ZA, Keen S, Teijlingen E Van. Risky work: Accidents among Nepalese migrant workers in Malaysia, Qatar and Saudi Arabia. Heal Prospect J Public Heal. 2017;3–10. doi: 10.3126/hprospect.v16i2.18643

7. Tripur Manandhar MV der P. Migrant Workers’ Psychological Wellbeing: The Case of Nepalese Construction Workers in Qatar. 2015. p. 140–6. Available from: https://s3.amazonaws.com/academia.edu.documents/38210948/Poceedings-New-Voices-2015.pdf?response-content-disposition=inline%3Bfilename%3DVulnerability_Shifts_in_a_Transitioning.pdf&X-Amz-Algorithm=AWS4-HMAC-SHA256&X-Amz-Credential=AKIAIWOWYYGZ2Y53UL3A%2F

8. Poudel OP, Thapa B, Bhandary S. Pre-departure psychological distress, depression, anxiety, stress and perception of risk factors at workplace among migrant workers of Nepal: A pilot study. J Gen Pract Emerg Med Nepal. 2019;(8):20–5. Available from: https://www.jgpeman.com

9. Lovibond, SH, Lovibond P. Manual for the depression anxiety stress scales. Psychol Found Aust. 1995. Available from: http://www2.psy.unsw.edu.au/dass//

10. Tonsing KN. Psychometric properties and validation of Nepali version of the Depression Anxiety Stress Scales (DASS-21). Asian J Psychiatr. 2018;8(December 2010):63–6. doi: 10.1016/j.ajp.2013.11.001

11. Saraswati LR, Rob U, Puri M, Sarna A. LIFE ACROSS THE BORDER: MIGRANTS IN SOUTH ASIA. 2015. Available from: https://assets.publishing.service.gov.uk/media/57a0897f40f0b649740000e8/61263_Final-Migrant-Report_Life-across-the-border.pdf

12. IOM Nepal. Research on the health vulnerabilities of the cross border migrants from Nepal. 2019. Available from: http://www.iom.int/nepal

13. Galappaththi R. Psychological Distress among Aspiring Sri Lankan Migrant Labour Workers and Associated demographic variables. 2018. Available from: https://www.academia.edu/38368829/

14. Ang JW, Chia C, Koh CJ, Chua BWB, Narayanaswamy S, Wijaya L, et al. Healthcareseeking behaviour, barriers and mental health of non-domestic migrant workers in Singapore. BMJ Glob Heal. 2017;2(2). doi: 10.1136/bmjgh-2016-000213

15. Lindert J, Priebe S, Mielck A, Bra E. Depression and anxiety in labor migrants and refugees-A systematic review and meta-analysis. Soc Sci Med. 2009;69:246–57. doi: https://doi.org/10.1016/j.socscimed.2009.04.032

16. Jasinskaja-lahti I, Yijälä A. The model of pre-acculturative stress—A pre-migration study of potential migrants from Russia to Finland. Int J Intercult Relations. 2011;35:499–510. doi: 10.1016/j.ijintrel.2010.11.003

17. Martínez-Ortega JM, Gutiérrez-Rojas L, Mendieta-Marichal Y, Jurado D, Gurpegui M, Alarcón RD. Factors associated with psychological distress or common mental disorders in migrant populations across the world. Rev Psiquiatr y Salud Ment (English Ed). 2017;10(1):45–58. doi: 10.1016/j.rpsmen.2017.02.004

18. Regmi PR, Aryal N, van Teijlingen E, Simkhada P, Adhikary P. Nepali Migrant Workers and the Need for Pre-departure Training on Mental Health: A Qualitative Study. J Immigr Minor Heal. 2019;(0123456789). doi: https://doi.org/10.1007/s10903-019-00960-z

19. Regmi PR, van Teijlingen E, Mahato P, Aryal N, Jadhav N, Simkhada P, et al. The Health of Nepali Migrants in India: A Qualitative Study of Lifestyles and Risks. Int J Environ Res Public Health. 2019;16(19):3655. doi: 10.3390/ijerph16193655

20. Simkhada P, Van Teijlingen E, Gurung M, Wasti SP. A survey of health problems of Nepalese female migrants workers in the Middle-East and Malaysia. BMC Int Health Hum Rights. 2018;18(1):1–7. doi: 10.1186/s12914-018-0145-7

